# Pooled CRISPR screening with single-cell transcriptome read-out

**DOI:** 10.1101/083774

**Authors:** Paul Datlinger, Christian Schmidl, André F Rendeiro, Peter Traxler, Johanna Klughammer, Linda Schuster, Christoph Bock

## Abstract

CRISPR-based genetic screens have revolutionized the search for new gene functions and biological mechanisms. However, widely used pooled screens are limited to simple read-outs of cell proliferation or the production of a selectable marker protein. Arrayed screens allow for more complex molecular read-outs such as transcriptome profiling, but they provide much lower throughput. Here we demonstrate CRISPR genome editing together with single-cell RNA sequencing as a new screening paradigm that combines key advantages of pooled and arrayed screens. This approach allowed us to link guide-RNA expression to the associated transcriptome responses in thousands of single cells using a straightforward and broadly applicable screening workflow.

## Main text

Pooled CRISPR screening is a powerful and widely used method for identifying critical genes involved in biological mechanisms such as cell proliferation^1, 2^, drug resistance^3^, and viral infection^4, 5^. Cells are infected in bulk with a library of guide-RNA (gRNA) encoding vectors, and the distribution of gRNAs is monitored before and after applying a selective challenge (Fig. 1a). Pooled CRISPR screens work well for mechanisms that affect cell survival and proliferation, and they can be extended to measure the activity of individual genes (e.g., by using engineered reporter cell lines). However, they cannot be combined with complex molecular read-outs such as transcriptome profiling, which is one of the most comprehensive and informative measures of cellular response^6^. Arrayed screens, in which only one gene is targeted at a time, make it possible to use RNA-seq as read-out^7^, but at the cost of a much lower throughput given that individual gRNAs have to be physically separated (Fig. 1b).

**Figure 1:**
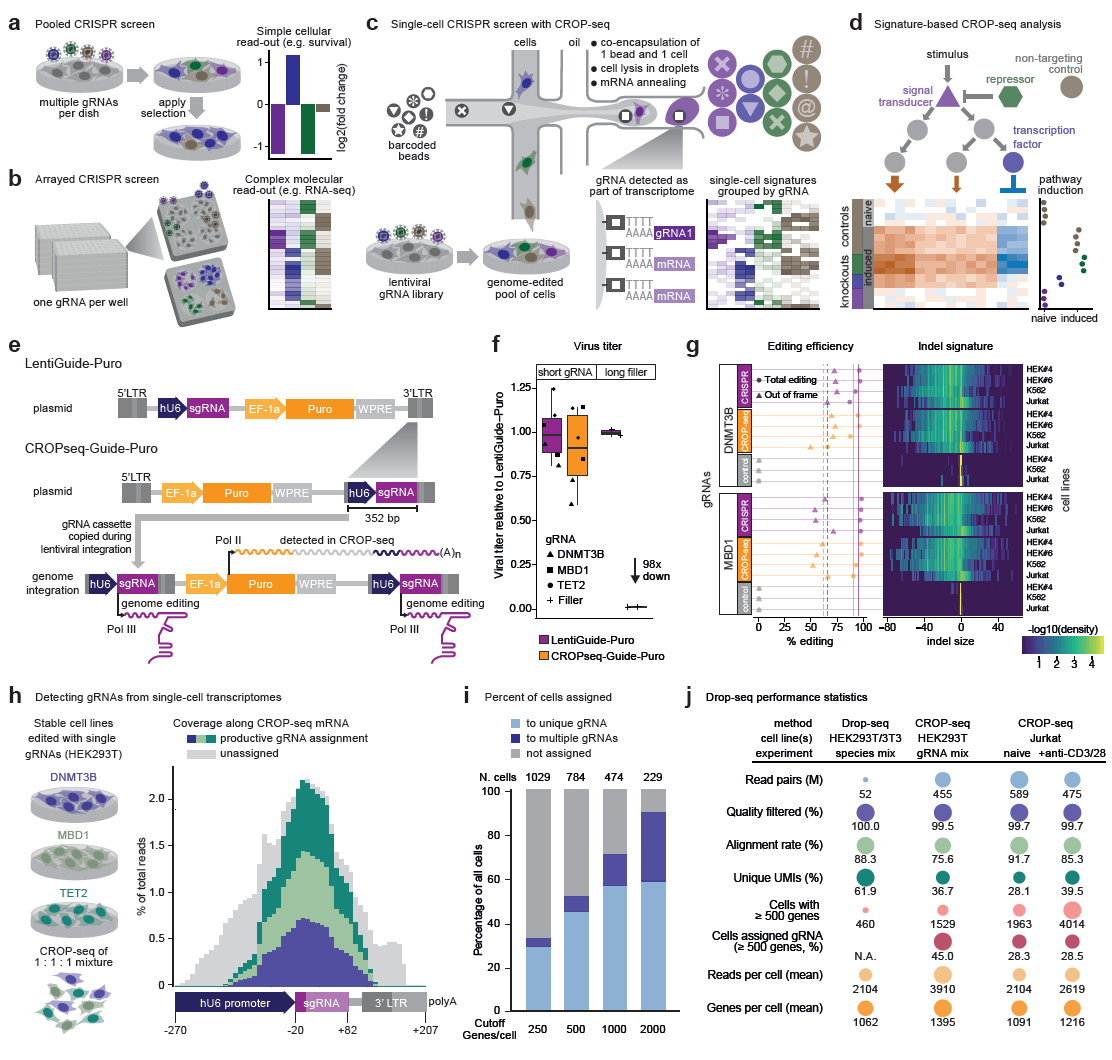
CROP-seq enables pooled CRISPR screening with single-cell transcriptome readout. **a)** Pooled screens detect changes in gRNA abundance in a bulk population of cells. They are limited to sim-ple read-outs such as cell proliferation, resistance to drugs or viruses, or expression of a selectable marker protein. b) Arrayed screens enable complex molecular read-outs such as transcriptome profiling. They typi-cally support only one gRNA per well. c) Single-cell CRISPR screens combine CRISPR genome editing in a bulk population of cells with a single-cell sequencing read-out. Specifically, the CROP-seq protocol uses a variation of Drop-seq to profile each cell’s transcriptome together with the expressed gRNA, and knockout signatures are derived by averaging across cells that express gRNAs for the same target gene. d) CROP-seq data analysis identifies pathway signature genes and quantifies the effect of specific gRNAs on these signa-tures. e) CROP-seq uses a lentiviral construct in which the gRNA expression cassette is positioned within the 3’ long-terminal repeat (LTR), causing its duplication during lentiviral integration. It is expressed as part of two transcripts – a short RNA polymerase III transcript for genome editing and a long RNA polymerase II transcript that is poly-ade-nylated and detectable with Drop-seq. **f)** Lentiviral titers for LentiGuide-Puro (a standard CRISPR vector) and the CROPseq-Guide-Puro vector show that cloning into the 3’ LTR did not compromise the lentiviral function for gRNAs (in contrast, a 1,885 bp filler construct led to a strong reduc-tion in viral titer). **g)** Genome editing efficiencies and indel signatures are similar between LentiGuide-Puro and CROPseq-Guide-Puro, based on next generation sequencing of PCR amplicons. **h)** CROP-seq can detect gRNAs from single-cell transcriptomes. **i)** Between 30% and 60% of single-cell transcriptomes are uniquely assigned to a gRNA, depending on the chosen threshold for the minimum number of genes detected in a cell. **j)** Performance statistics for single-cell RNA-seq across all CROP-seq experiments.

Here we propose a third and complementary screening paradigm, single-cell CRISPR screens, based on the idea that gRNAs and their cellular response are already compartmentalized within single cells. Detecting each cell’s gRNA along with the corresponding single-cell transcriptome thus enables the derivation of gene expression signatures for individual gene knockouts in a complex pool of cells (Fig. 1c). Our CRISPR Drop-seq (CROP-seq) method comprises a gRNA vector that makes individual gRNAs detectable in single-cell RNA-seq experiments, a high-throughput assay for single-cell RNA-seq^8^, a computational pipeline for assigning single-cell transcriptomes to gRNAs, and a bioinformatic method for analyzing and interpreting gRNA-induced transcriptional profiles. Combining these building blocks, CROP-seq enables pooled screens with single-cell transcriptome read-out, for example for dissecting complex signaling pathways that are not easily reduced to a single selectable marker (Fig. 1d). gRNAs are typically transcribed by RNA polymerase III from a human U6 promoter (hU6), hence they lack a polyadenylated tail and are not detectable with most single-cell RNA-seq protocols. We thus redesigned a popular construct for pooled CRISPR screening (LentiGuide-Puro)^9^ to produce the gRNA both in its functional form and as part of a polyadenylated mRNA transcript (Fig. 1e). By incorporating the hU6-gRNA cassette into the lentiviral 3’ LTR, it becomes part of the puromycin resistance mRNA transcribed by RNA polymerase II and detectable by RNA-seq protocols based on poly-A enrichment, while the functional gRNA continues to be expressed from the hU6 promoter. As most single-cell RNA-seq technologies focus on the 3’-end of mRNAs, we placed the hU6-gRNA cassette as close to the AATAAA polyadenylation motif as possible to reproducibly recover gRNA sequences from the transcriptome. The entire hU6-gRNA cassette is also duplicated along with the 3’ LTR during reverse transcription and integration of the virus (Supplementary Fig. 1), which results in a second copy upstream of the possibly interfering EF-1a promoter – a design reminiscent of an early shRNA expression vector^10^.

Having assembled our CROPseq-Guide-Puro plasmid using ligase cycling reaction^11^, we validated its performance and suitability for CRISPR screening in a series of experiments. We found that the hU6-gRNA insertion into the 3’ LTR had no adverse effects on the formation of lentiviral particles nor on the stability of the puromycin mRNA (Fig. 1f). We produced virus from LentiGuide-Puro and CROPseq-Guide-Puro containing either one of three validated gRNAs (targeting DNMT3B, MBD1, or TET2) or a longer filler construct that was expected to interfere with viral activity (Fig. 1f), and we observed comparable functional titers for all gRNA sequences tested. Changes to the construct are therefore well-tolerated, as long as the size of the 3’ LTR insertion stays within certain limits, as exemplified by the massive reduction in viral titer observed for the 1,885 bp filler.

The genome editing efficiencies of LentiGuide-Puro and CROPseq-Guide-Puro were systematically compared in three Cas9-expressing cell lines (K562, Jurkat, and two clones of HEK293T) with two gRNAs and two complementary technologies: the T7 endonuclease assay^12^ (Supplementary Fig. 2) and next generation sequencing of PCR amplicons (Fig. 1g). Genome editing was very efficient for both constructs, averaging at 95.5% (LentiGuide-Puro) and 90.5% (CROPseq-Guide-Puro), with the expected two thirds (66.2% and 62.4%) of molecules carrying out-of-frame insertions and deletions. Using the sequencing reads to define the exact length of the induced indels, we observed that different cell lines and genomic loci exhibit characteristic editing signatures (e.g. pronounced insertions in Jurkat cells), which were highly similar for both constructs (Fig. 1g). We thus concluded that CROPseq-Guide-Puro yields excellent results both in terms of the quantity and quality of induced genome editing events.

Next, we tested whether we could detect expressed gRNAs in single-cell transcriptome data obtained for CROPseq-Guide-Puro infected cells using a variation of the Drop-seq protocol^8^ (Fig. 1h and Supplementary Fig. 3). We generated three stable knockout cell lines from HEK293T cells, targeting either DNMT3B, MBD1, or TET2 with a single gRNA. By performing the infections independently, we ensured that each cell received only one gRNA – even in the rare case of multiple infection events. Performing Drop-seq on a 1:1:1 mixture of the three cell lines, we observed high sequencing coverage at the gRNA (Fig. 1h), which allowed us to assign between 30% and 60% of high-quality transcriptomes to one gRNA (Fig. 1i). Based on this mixing experiment, a quality threshold of at least 500 detected genes per cell was selected, resulting in 45% uniquely assigned cells. We also estimated the rate of false gRNA assignments based on the number of cells with more than one gRNA, which should not happen in our experimental setup. At our chosen quality threshold, around 12.5% (98 out of 784) of assigned cells fall into this category (Fig. 1i). While significant, these false assignments can be expected to average out when aggregating data across multiple single cells as described below.

Having established and validated CROP-seq as a method for single-cell CRISPR screens (see Fig. 1j for performance statistics of the presented CROP-seq experiments), we applied our method in a biologically motivated screen of T cell receptor (TCR) activation in Jurkat cells (Fig. 2a). We designed a gRNA library for six high-level regulators of TCR signaling and 23 transcription factors that may act further downstream, targeting each gene with three distinct gRNAs. Furthermore, we added 20 previously published non-targeting gRNAs as negative controls and nine gRNAs targeting essential genes^2^ as positive controls. Jurkat cells that stably express Cas9 were transduced with the CROPseq-Guide-Puro lentiviral library at a low multiplicity of infection, and gRNA expressing cells were enriched by puromycin selection. At day 10 post infection, the surviving pool of genome-edited cells was serum-starved, split, and subjected either to TCR stimulation via CD3 and CD28 or to continued starvation, and both conditions were analyzed with CROP-seq.

**Figure 2:**
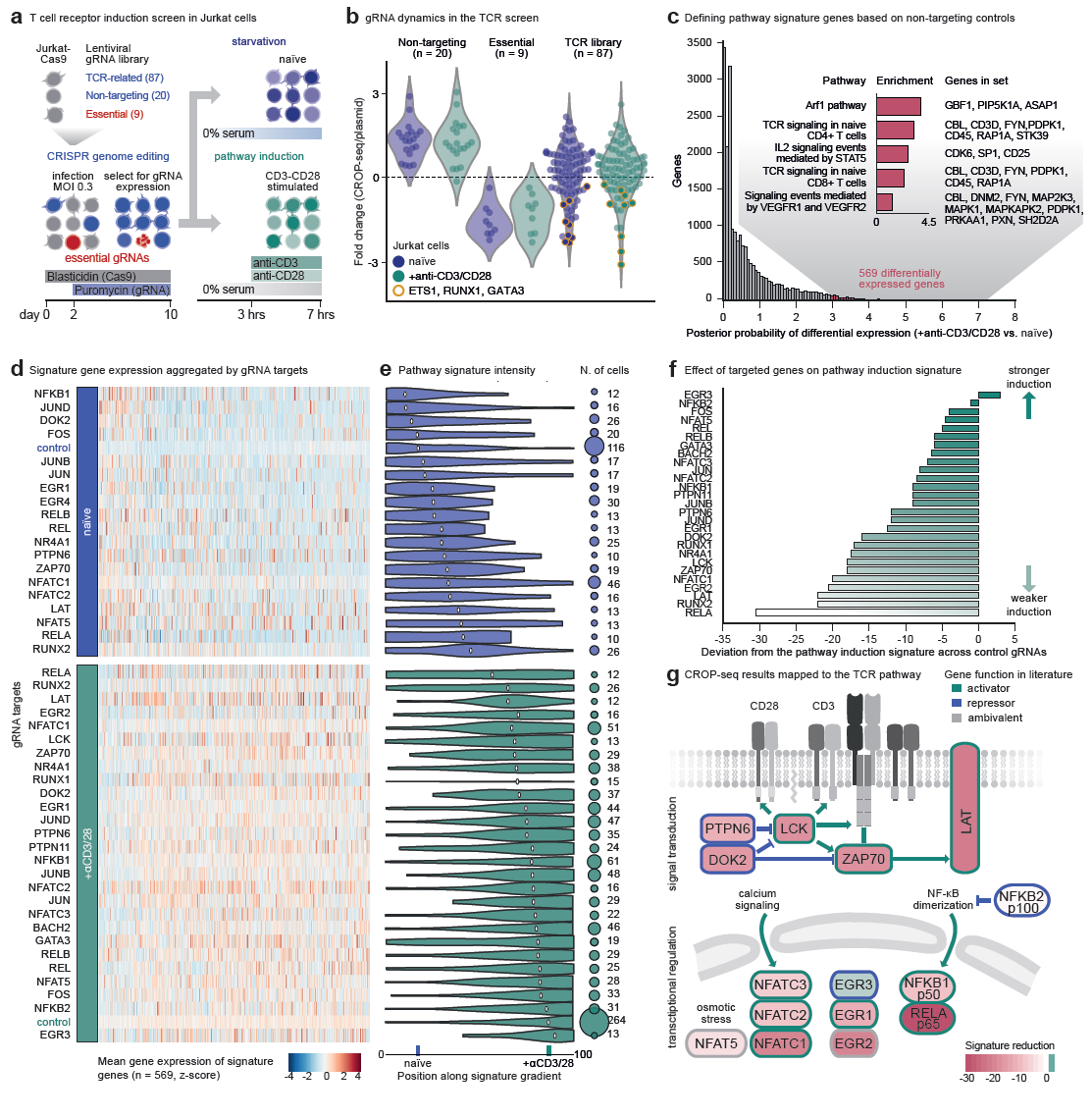
CROP-seq analysis of T cell receptor signaling. **a)** Experimental design of a single-cell CRISPR screen for the T cell receptor (TCR) pathway. Cas9-expressing Jurkat cells were infected with the CROPseq-Guide-Puro lentivirus containing a TCR-focused gRNA library at a low multiplicity of infection (MOI = 0.3). Lentivirus expressing cells were selected, and after 10 days the cell population was serum-starved, split, and either stimulated with anti-CD3/CD28 anti-bodies, or subjected to continued starvation. Cells were then subjected to single-cell RNA-seq. b) Fold change of gRNA abundance between the original plasmid pool and the CROP-seq experiments for naïve and TCR-stimulated cells. Values were normalized to the total of detected amplicons or assigned cells, re-spectively. c) Differential expression between TCR-stimulated cells and naïve Jurkat cells (histogram) and enriched pathways among the most differentially expressed genes (inlet). The x-axis shows the uncorrected posterior probability, and the chosen threshold for differentially expressed genes (red) corresponds to a false discovery rate of 5% using Holm-Bonferroni correction25. The five most enriched pathways based on the Enrichr tool26 tool are shown and sorted by their combined enrichment score (p-value * z-score). d) Heatmap of mean relative expression (column z-score) across the 569 pathway signature genes (columns), aggregated across cells that express gRNAs targeting the same gene (rows). e) Distribution of signature gene intensity across single cells (left) and number of cells (right) grouped by gRNA target gene. (f) Deviation of signature gene intensity for each gRNA target gene relative to control cells in TCR-stimulated Jurkat cells. g) CROP-seq results (signature gene intensity as shown in panel f) mapped to key components of the TCR pathway.

To monitor gRNA dynamics over the course of the screen, we compared the gRNA assignments based on CROP-seq to gRNA counts obtained by sequencing the plasmid library. Using this metric, we observed consistent depletion of gRNAs targeting essential genes (positive controls), which validates the editing efficiency of our screen (Fig. 2b). The gRNAs in our TCR library showed more diverse patterns: gRNAs targeting the transcription factors ETS1, RUNX1, and GATA3 were depleted in the same way as the positive controls, suggesting that they are essential in Jurkat cells. In contrast, most gRNAs showed patterns that were more similar to the negative controls, indicating no strong anti-proliferation effect.

To establish a reference for the transcriptome response to induced TCR signaling, we combined the transcriptome data from all single cells that were assigned to the non-targeting gRNAs (negative controls). Comparing the expression levels between anti-CD3/CD28-stimulated and naïve Jurkat cells, we identified a set of 569 differentially expressed genes, which constitute a pathway signature of the TCR response in our model. These genes were enriched for gene sets relevant to TCR signaling (Fig. 2c; Supplementary Fig. 5 and 6), which provides external validation of the validity of the cellular system and of the single-cell RNA-seq data quality.

Next, we aggregated the single-cell expression profiles by the target genes of the detected gRNAs (Fig. 2d) and quantified the strength of the pathway signature in each single cell. The resulting distribution of single-cell pathway signature intensities placed each cell on a gradient between the naïve and stimulated control Jurkat cells (Fig. 2e). We then calculated the deviation of the mean for each target gene from the mean of the control cells under the same conditions (Fig. 2f and Supplementary Fig. 7), which captures the induced shift in pathway activity when the target gene is inactivated. This analysis was based on 1,612 high-quality single cell gene expression profiles with unique gRNA assignments, 10 to 61 cells per targeted gene, and 380 control cells used as reference.

Focusing on the stimulated cells, we observed that gRNAs for target genes immediately downstream of the TCR (such as the kinases LCK and ZAP70 and the adapter protein LAT) had a strong negative effect on the TCR activation signature, consistent with their crucial role in signal propagation and amplification^13^ (Fig. 2g). Accordingly, targeting key transcription factors downstream of TCR signaling from the NF-κB^14^, NFAT^15^, and EGR^16^ families (e.g. RELA, NFATC1, and EGR1) strongly decreased the TCR activation signature. In contrast, transcription factors that were not directly related to TCR signaling (e.g., NFATC5, which plays a role in osmotic stress induced lymphocyte activation^17^) had little effect. Finally, among the known negative regulators of T cell activation, we found that only EGR3-targeted cells displayed a slightly increased TCR induction signature in our model.

In addition to studying known regulators of TCR response as a validation of our method, we explored whether CROP-seq can help understand the role of transcription factors with unresolved biological function. For example, the targeting of RUNX2, NR4A1, and EGR2 strongly decreased the TCR activation signature in our model, indicating a potential role of these proteins in TCR signaling. This is in line with recent reports identifying RUNX2 as a potential activator of an early CD8 cytokine and effector signature^18^ and with the surprising finding that EGR2 can have activating functions during the differentiation of naïve peripheral T cells and influenza infection^19^.

Through this proof-of-concept screen, we have demonstrated the feasibility of single-cell CRISPR screening, and we established CROP-seq as a method for pooled CRISPR screening with single-cell transcriptome sequencing read-out. Importantly, we showed that relevant transcriptome signatures can be derived directly from the single cell data, which makes prior knowledge of the expected expression changes dispensable. We expect CROP-seq to be broadly useful for studying complex biological phenomena that are difficult to reduce to a simple read-out amenable to classical pooled screens. Moreover, with increasing throughput of single-cell transcriptomics^20^ and the advent of single-cell multi-omics technology^21^, CROP-seq has the potential to provide comprehensive characterization of large CRISPR libraries and a powerful method for dissecting cellular regulation at scale.

## Methods

### Cloning and validation of the CROPseq-Guide-Puro plasmid

To clone CROPseq-Guide-Puro, we amplified PCR products from LentiGuide-Puro (Addgene #52963) and assembled them in different order using the Ligase Cycling Reaction (LCR)^11^. We screened for correctly assembled clones by colony PCR and further validated the assembly by restriction digestion with either SphI or SapI as well as Sanger sequencing. Moreover, to validate the duplication of the hU6-gRNA cassette, genomic DNA from cells infected with LentiGuide-Puro or CROPseq-Guide-Puro lentivirus, or the CROPseq-Guide-Puro plasmid pool were PCR-amplified. Productive amplification occurs only when the gRNA cassette is duplicated during lentiviral reverse transcription and integration or when amplifying from a circular plasmid.

### Cloning of individual gRNAs into the CROPseq-Guide-Puro plasmid

gRNA cassettes were annealed from two oligos (top: 5’-CACCG(N)_20_-3’, bottom: 5’-AAAC(N)_20_C-3’) by combining 1 μl of each 100 μM oligo with 1 μl of 10x T4 ligation buffer (NEB #B0202S), 6.5 μl of water, and 0.5 μl of T4 Polynucleotide Kinase (NEB #M0201S), incubating as follows: 37 °C for 30 min (oligo phosphorylation), 95 °C for 5 min, then ramping from 90 °C to 25 °C at 5 °C/min. Plasmid backbone was prepared by digesting 1 μg of CROPseq-Guide-Puro with 10 units of BsmBI (NEB #R0580L) in a volume of 30 μl 1x NEB buffer 3.1, incubating for 1 hour at 55 °C. To dephosphorylate the digested plasmid, we added 2 μl of Shrimp Alkaline Phosphatase (rSAP, NEB #M0371L), incubating for 1 hour at 37 °C followed by heat-inactivation (both BsmBI and rSAP) for 20 min at 80 °C. Ligation reactions were set up as follows: 1.6 μl of rSAP reaction, 1 μl gRNA cassette (diluted 1:200 in water), 5 μl 2x Quick Ligase buffer, 2.4 μl water, and 1 μl Quick Ligase (NEB #M200S), incubated at 25 °C for 15 min. The ligation reaction was chemically transformed into the NEB stable *E. coli* (NEB #C3040H) following the manufacturer’s high efficiency protocol.

### Amplification-free cloning of pooled gRNA libraries into the CROPseq-Guide-Puro plasmid

Vector backbone was prepared by digesting 5 μg of CROPseq-Guide-Puro with 20 U of BsmBI (NEB #R0580L) in a total volume of 25 μl 1x NEB buffer 3.1, incubating for 1 hour at 55 °C and inactivating the restriction enzyme for 20 min at 80 °C. The 8,332 bp fragment was purified using the SNAP UV-free gel purification kit (Invitrogen #K200025). gRNAs were synthesized by Sigma Aldrich as 74 base oligos, with 18 and 35 bases of homology to the hU6 promoter and guide RNA backbone, respectively. Oligos were diluted to 100 μM and pooled in equal amounts. The oligo pool was further diluted 1:1000 for a final concentration of 0.1 pg/μl for assembly into the vector backbone. gRNA libraries were cloned by Gibson’s isothermal assembly: 11 fmoles (56.7 ng) of CROPseq-Guide-Puro backbone and 200 fmoles (0.2 pg) of pooled ssDNA oligos were combined with 10 μl of NEBuilder HiFi DNA assembly master mix and water to 20 μl. After 1 hour of incubation at 50 °C, reactions were desalted by filter dialysis (Merck #VMWP04700) and 10 μl of the reaction were electroporated into 25 μl of Lucigen Endura *E. coli* cells (Lucigen #60242-2) using pre-chilled 1 mm electroporation cuvettes (BioRad #1652089) in a BioRad GenePulser I machine set to 25 μF, 200 Ω and 1.5 kV. Within seconds after the pulse, 1 ml of 37 °C Recovery Medium (Lucigen) was added and bacteria were grown in round-bottom tubes for 1 hour at 37 °C while shaking at 180 rpm. Then, 1 ml of the bacterial culture was plated on a 25x25 cm bioassay plate containing LB medium (Miller) with 100 μg/ml carbenicillin. Plates were incubated at 32 °C for 22 hours, then LB medium was added and cells were scraped off the plate. Bacterial cells were pelleted by 15 min of centrifugation at 5000 rcf at 4 °C and plasmid DNA was extracted with Qiagen’s EndoFree Plasmid Mega kit (Qiagen #12381). Library coverage was estimated by counting the number of bacterial colonies on a 1:1000 dilution plate. The T cell receptor library was cloned at 853x coverage.

### Lentivirus production

HEK293T cells were plated onto 6-well plates at 1.2 million cells/well in 2 ml of lentivirus packaging medium [Opti-MEM I (Gibco #51985-034), 5% FCS (Sigma), 200 μM sodium pyruvate (Gibco #11360-070), no antibiotics]. The next morning, cells were transfected with lipofectamine 3000 (Invitrogen #L3000015) using 1.7 μg of CROPseq-Guide-Puro (containing single gRNAs or libraries) or lentiCas9-Blast (Addgene #52962), and 0.9 μg each of the three packaging plasmids pMDLg/pRRE (Addgene #12251), pRSV-Rev (Addgene #12253) and pMD2.G (Addgene #12259). The medium was exchanged for fresh lentivirus packaging medium 6 hours after the transfection. Supernatant containing viral particles was harvested at 24 and 48 hours and passed through a 0.45 μm filter to remove cells. Further concentration of viral particles was not required. Virus was stored at −80 °C, later one vial was thawed for estimating the virus titer.

### Lentivirus titration

HEK293T cells were seeded onto 24-well plates at 5×10^4^ cells/well in 500 μl of culture medium [DMEM (Gibco #10569010), 10% FCS, no antibiotics] and grown over night to reach 30% to 50% confluence. The next day, the medium was exchanged for 450 μl/well of fresh culture medium (prepared as described above) containing 8 μg/ml polybrene (Sigma Aldrich #H9268-5G), which was also used to dilute the viral stock. Lentivirus preparations were thawed from storage at −80 °C, and titrated in a 1:5 dilution series ranging over 10 wells (1:5 to 1:9,765,625). Each dilution was tested in duplicate, by adding 50 μl/well to the 24-well plate. At least two wells per plate served as untransduced controls. 24 hours after the infection, the medium was exchanged for 500 μl/well of complete culture medium [DMEM (Gibco #10569010), 10% FCS, penicillin/streptomycin]. Starting at 48 hours post infection, the medium was exchanged for selection medium every 2-3 days. Depending on the construct, the selection medium for HEK293T cells contained either 22.5 μg/ml of blasticidin (Invivogen #ant-bl-5) or 2.25 μg/ml of puromycin (Fisher Scientific #A1113803). As soon as all cells in the untransduced controls had died, medium was removed and the plate was washed once with 1x PBS. Resistant colonies were stained for 15 min in a solution of 1% (w/v) crystal violet (Sigma Aldrich #61135-25G) in a 10% solution of ethanol in water. After washing the plate several more times with 1x PBS, colonies were counted and the virus titer (transducing units/ml) was calculated as [(# of resistant colonies) x dilution factor] * (1000 μl/50 μl).

### Lentiviral transduction with gRNA libraries or single gRNAs

For suspension cell lines (Jurkat), cells were seeded in 6-well plates at 5 million cells/well in 2 ml of complete culture medium [RPMI (Gibco #21875-034) with penicillin/streptomycin and 10% FCS (Sigma)] containing 8 μg/ml polybrene (Sigma Aldrich #H9268-5G) and blasticidin (Invivogen #ant-bl-5) to keep up selection for lenti-Cas9-Blast. Wells 1-5 were infected with 250 μl and 50 μl of the virus stock and 50 μl of 1:3, 1:6 and 1:12 dilutions. Well 6 served as the uninfected control. Immediately after addition of the virus, cells were centrifuged at 37 °C and 1200 rcf for 45 min, followed by overnight incubation at 37 °C, 5% CO_2_. After 24 hours, cells were pelleted in 15 ml tubes, taken up in 30 ml of fresh complete culture medium containing blasticidin and transferred to T75 culture flasks. At 48 hours post infection, selection with puromycin (Fisher Scientific #A1113803) was started. A flask with about 30% surviving cells was chosen and grown for 10 days to allow for efficient genome editing, while renewing the selective medium (containing blasticidin and puromycin) every 2-3 days. Adherent cells (HEK293T) were plated at 5 million cells per 15 cm dish in complete culture medium [DMEM (Gibco #41965-039), 10% FCS, penicillin/streptomycin], preparing six plates per library. After 24 hours, the medium was replaced by complete culture medium containing 8 μg/ml polybrene and different amounts of virus were added: 250 or 50 μl of the stock, or 50 μl of a 1:3, 1:6 or 1:12 dilution. The sixth plate remained uninfected. After 24 hours, the virus-containing medium was replaced by fresh complete culture medium. Puromycin selection was started at 48 hours post infection and 2 days later, a plate with about 30% surviving cells was selected and grown under blasticidin and puromycin selection for 10 days.

### T7 endonuclease assay

For easy-to-resolve cleavage products, PCR primers were designed such that the DNMT3B and MBD1 gRNA targets are located off-center of the amplicons. PCR reactions were set up in a reaction volume of 50 μl: 25 μl of Q5 Hot Start High-Fidelity 2x Master Mix (NEB #M0494L), 2.5 μl 10 μM FWD primer, 2.5 μl 10 μM REV primer, 100 ng of genomic DNA and water, and incubated as follows: 98 °C for 30 s, 35x [98 °C for 5 s, Ta for 10 s, 72 °C for 20 s], 72 °C for 2 min and storage at 10 °C. PCR products were purified by a 2.0x AMPure XP bead clean-up (Beckman Coulter #A63880) and measured in a Qubit HS assay (Invitrogen #Q32854). 200 ng of each PCR product were taken up in 19 μl of 1x NEB buffer 2 and subjected to denaturation and re-annealing: 95 °C for 5 min, 95 to 85 °C at −2 °C/s, 85 to 25 °C at −0.1 °C/s, hold at 4 °C. Mismatched DNA duplexes were then digested by addition of 1 μl T7 endonuclease I (NEB #M0302) followed by incubation at 37 °C for 15 min. The reaction was stopped with 1.5 μl of 0.25 M EDTA (Invitrogen #15575020) and 1 μl was analyzed on an Agilent High Sensitivity DNA chip (Agilent #5067-4616), yielding the molarity of digested and undigested fragments from which the editing efficiencies were calculated as: average(digested1, digested2) / sum(average(digested1, digested2), undigested).

### Indel analysis by next generation sequencing

Amplicons were designed to be smaller than 500 bp, with the gRNA target site at the center. PCR was performed as described for the T7 endonuclease assay. PCR products were purified by a 2.0x AMPure XP bead cleanup (Beckman Coulter #A63880) and used as input for Nextera XT (Illumina #15032350) library preparations, according to the standard protocol. Libraries were sequenced on the Illumina MiSeq platform using a 150-cycle v3 flow cell with dual indexing. The machine was set to read lengths of 159 (read1) + 8 (i7) + 8 (i5) bases. To analyze the data, we first defined two 10 bp-long ’anchor’ sequences on both sides of the gRNA at a fixed distance of 30 bp. We then extracted reads spanning the gRNA target site from the BAM file via a grep operation for the pattern “<anchor_left>.*<anchor_right>” on the BAM file, using the -o option to return only the matching part of the sequence. For wild type fragments, this sequence is exactly 100 bp long: 10 bp anchor_left + 30 bp + 20 bp (gRNA) + 30 bp + 10 bp anchor_right. The size of insertions and deletions was calculated as the deviation from the wild type length, summarized, and plotted.

### Testing plasmid library complexity and gRNA dynamics in the pooled screen

The gRNA expression cassette was amplified from either the plasmid library or the genomic DNA of cells at day 10 post infection. A single PCR reaction is sufficient to introduce all functional sequences required for compatibility with Illumina machines. We included 8 bp long i7 barcodes and a stagger sequence to increase library complexity, resulting in a set of variations of primers that can be freely combined. Purified PCR products were pooled and sequenced on an Illumina MiSeq machine, using a 150-cycle v3 flow cell with a read configuration of 159 (read1) + 8 (i7) + 8 (i5). Read1 covers the stagger sequence (of variable length), the FWD primer binding site (23 bases), the gRNA target (20 bases), and the backbone (82 bases). Guide RNA read counts were normalized by the total number of reads sequenced. Counts of cells assigned gRNAs (regardless of the size of the captured transcriptome per cell) were normalized to the total number of assigned cells. Fold change values were calculated between normalized cell counts from the CROP-seq screen and the normalized gRNA counts from the plasmid library.

### CD3/CD28 stimulation of the T cell receptor pathway in Jurkat cells

Jurkat cells were serum-starved for 3 hours prior to stimulation. They were stimulated with 1 μg/ml anti-CD3 (eBioscience #16-0037) and 1 μg/ml anti-CD28 antibody (eBioscience #16-0289-81) for 4 hours while under starvation and then subjected to CROP-seq. The unstimulated (naïve) control was subjected to continued starvation.

### Drop-seq protocol for highly multiplexed single-cell RNA-seq in microfluidic droplets

Adherent cells were detached using Trypsin-EDTA (Gibco #25300-054), following standard cell culture practices. Cells were collected by centrifugation at 300 rcf for 5 min, washed once with PBS-0.01% BSA (freshly prepared on the day of the run), and resuspended in 1 ml of PBS-0.01% BSA. Cells were filtered through a 40 μm cell strainer to obtain a suspension of single-cells, which were counted using a CASY device. Single cells were then co-encapsulated with barcoded beads (ChemGenes #Macosko-2011-10) using an Aquapel-coated PDMS microfluidic device (FlowJEM), connected to syringe pumps (kdScientific) via polyethylene tubing with an inner diameter of 0.38 mm (Scicominc #BB31695-PE/2). Cells were supplied in PBS-0.01% BSA at a concentration of 220 cells/μl, barcoded beads were resuspended in Drop-seq lysis buffer at a concentration of 150 beads/μl. The flow rates for cells and beads were set to 1.6 ml/hour, while Droplet Generation Oil (BioRad #1864006) was run at 8 ml/hour. During the run, the barcoded bead solution was mixed by magnetic stirring with a mixing disk set to 1 jump/s. A typical run lasted between 35 and 40 min. In case of multiple runs per day, droplets were intermittently stored at 4 °C and processed together. Our most important modification to the protocol is an alternative way to break droplets, which recovers beads much more efficiently than in the original publication of Drop-seq^8^. After removing as much oil below the droplet layer as possible, we added 30 ml of 6x SSC buffer (Promega #V4261) and 1 ml of Perfluoroctanol (Sigma Aldrich #370533-25G) and shook the tube forcefully 6 times to break the droplets. Based on their large diameter, beads were then collected by syringe-filtering the solution through a 0.22 μm filter unit (Merck #SLGV033RS), washing 2x with 20 ml of 6x SSC buffer and eluting by turning the filter upside down and rinsing it with 3x 10 ml of 6x SSC buffer. Beads were then collected by centrifugation at 1,250 rcf for 2 min, setting the brake speed to 50%. After washing a second time with 10 ml 6x SSC, the pellet was taken up in 200 μl of 5x RT buffer and transferred to a 1.5 ml tube. Reverse transcription and Exonuclease I treatment were performed as described in the original publication^8^, and the number of barcoded beads was estimated using a Fuchs-Rosenthal counting chamber (mixing the bead suspension with 6x DNA loading dye). Depending on the performance of the experiment, we prepared up to 24 PCR reactions per Drop-seq run, adding 4,400 beads (~220 cells) per PCR and enriching the cDNA for 4 + 13 cycles, using the already described reagents. We then prepared Drop-seq libraries using the Nextera XT kit (Illumina #15032350), starting from 1.5 ng of cDNA pooled in equal amounts from all PCR reactions for a given run. We typically required an additional 11 enrichment cycles, using the Illumina Nextera XT i7 primers along with the Drop-seq New-P5 SMART-PCR hybrid oligo. The slightly increased cDNA input typically results in an average size distribution of about 575 bp. After quality control, libraries were sequenced with paired-end SBS chemistry on Illumina HiSeq 3000/4000 instruments, leveraging the patterned flow cell technology. Drop-seq Custom Read1 Primer was spiked into the HP10 primer solution, located in column 11 of the cBot Reagent Plate at 1μM final concentration. High sequence complexity needed for optimal base calling performance was achieved by adding 20-30% PhiX as spike-in. Cluster generation and Read 1 primer hybridization were completed using cBot protocol ‘HiSeq_3000_4000_HD_Exclusion_Amp_v1.0’.

### Preprocessing of single-cell sequencing data

Single-cell sequencing data were processed using the Drop-seq Tools v1.12 software^8^. Briefly, each transcriptome Read 2 was tagged with the cell barcode (bases 1 to 12) and UMI (unique molecular identifier) barcode (bases 13 to 20) obtained from Read 1, trimmed for sequencing adapters and poly-A sequences, and aligned using STAR v2.4.0^22^ to a concatenation of the mouse and human genomes (for the species mixing experiment) or to the human reference genome assembly (Ensembl GRCm38 release) containing artificial chromosomes that represent the CROPseq-Guide-Puro plasmid construct. Reads aligning to exons were tagged with the respective gene name, and counts of unique UMIs per gene within each cell were used to build a digital gene expression matrix for cells with counts in at least 500 genes. Normalization of the expression matrix was performed with the R package Seurat^23^, first with log-transformation and scaling each cell to a total of one thousand molecules, including only genes present in at least 1% of all cells and excluding cells with more than the 95^th^ percentile of mitochondrial gene content across all cells. For the assignment of gRNAs to cells, we quantified the overlap of reads to the specific gRNA sequence within the CROPseq-Guide-Puro plasmid chromosomes and assigned the most abundant gRNA to the respective cell. For the analysis shown in Fig. 1i, cells were classified as uniquely assigned if the sequencing data had at least three times more overlap with the dominant gRNA than the sum of all other gRNAs; they were classified as containing multiple gRNAs where this difference was smaller than three; and they were classified as unassigned where no reads were overlapping with the gRNA sequence.

### Transcriptome signature analysis

Sparse transcriptome signatures were used to position single cells sharing a putative perturbation (in this case, gene knockouts as determined by the gRNA assignment) within a multidimensional axis of cell states (here, the measured activation of the TCR pathway in Jurkat cells). This method is reminiscent of the method for assessment of cross-contamination between cell types in single cells described previously^24^. We identified genes associated with TCR response (signature genes), selected cells assigned to a non-targeting control gRNA as representatives of the unperturbed stimulation (control cells), produced *in silico* mixtures of gene expression profiles of increasingly more stimulated cells based on the control cells from both conditions (mix profiles), and identified the mixed profile that best matched the expression profile of each single cell (signature position).

To determine a TCR-specific activation signature we used only cells containing a non-targeting control gRNA, and we compared the read counts of naïve cells with cells under the TCR activating CD3/CD28 stimulation condition using the single-cell differential analysis R package SCDE^25^, from which we developed dropout error models independently for each stimulation condition. Signature genes were defined by having an absolute, multi-testing corrected, posterior probability higher than 1.5. Bioinformatic analysis of the signature gene function was performed with the gene list enrichment analysis tool Enrichr^26^ using the rank of differential expression posterior as weight; the retrieved combined score (log[*p*-value] * *z*-score) was displayed.

Based on the mean expression level of cells assigned to non-targeting control gRNAs, for the signature genes we constructed a matrix *Z* of stepwise weighted linear mixtures of expression profiles between the unstimulated and stimulated conditions, which is given by:

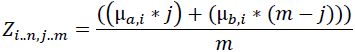

where μ_*a*_ and μ_*b*_ are vectors with mean expression values of cells in the unstimulated and stimulated conditions respectively, and *i* and *j* are indexes of the signature genes (*n*) and the number of desired mixtures *m*. We selected *m* = 100, and to account for cell-to-cell variability as well as overshooting changes in the signature genes compared to the mean of the control cells, we generated an extended linear space of length *m* with boundaries −20 and 120. For *m* = 100, this is equivalent to μ_*a*_ and μ_*b*_ being placed at index *j* = 20 and *j* = 80 of the *Z* matrix, respectively.

We positioned each single cell in the matrix by retrieving the argmax of the Pearson correlation of each cell to each mixture in the matrix. By grouping cells by their gRNA assignment, we visualized the distribution of cell signature positions per group and calculated the mean group signature position for groups with more than 10 cells. We then calculated the log fold deviation of each group of cells to the group of control cells within the respective stimulation condition, to determine a relative deviation group signature positions relative to genetically unperturbed cells.

### Reagent and data availability

The CROPseq-Guide-Puro plasmid will be made available via Addgene. A step-by-step experimental protocol for CROP-seq is being prepared and will be made available via a supplementary website. All raw and processed data will be made available via a supplementary website and via the NCBI GEO database.

## Acknowledgements

We would like to thank Astrid Fauster, Johannes Bigenzahn, and Michel Owusu for kindly providing Cas9 expressing cell lines, Matthias Farlik and Thomas Krausgruber for contributing to the Drop-seq setup, the Biomedical Sequencing Facility at CeMM for assistance with next generation sequencing, and all members of the Bock lab for their help and advice. CRISPR/Cas9 plasmids lentiCas9-Blast and lentiGuide-Puro were provide by Feng Zhang via Addgene (Plasmid #52962, #52963). Lentiviral packaging plasmids pMDLg/pRRE, pRSV-Rev and pMD2.G were provided by Didier Trono by Addgene (Plasmid #12251, #12253, #12259). C.S. is supported by a Feodor Lynen Fellowship of the Alexander von Humboldt Foundation. C.B. is supported by a New Frontiers Group award of the Austrian Academy of Sciences and by an ERC Starting Grant (European Union’s Horizon 2020 research and innovation programme, grant agreement n° 679146).

## Author contributions

Conceptualization, P.D., C.S., and C.B.; methodology, P.D., C.S., A.R., and C.B.; formal analysis, P.D., A.R., and J.K.; investigation, P.D., C.S., P.T., and L.S.; writing – original draft, P.D, C.S., A.R., and C.B.; writing – review & editing, P.T., J.K., and L.S.; supervision, C.B.; funding acquisition, C.B.

## Figure legends

**Figure 1: CROP-seq enables pooled CRISPR screening with single-cell transcriptome readout**

**a)** Pooled screens detect changes in gRNA abundance in a bulk population of cells. They are limited to simple read-outs such as cell proliferation, resistance to drugs or viruses, or expression of a selectable marker protein. **b)** Arrayed screens enable complex molecular read-outs such as transcriptome profiling. They typically support only one gRNA per well. **c)** Single-cell CRISPR screens combine CRISPR genome editing in a bulk population of cells with a single-cell sequencing read-out. Specifically, the CROP-seq protocol uses a variation of Drop-seq to profile each cell’s transcriptome together with the expressed gRNA, and knockout signatures are derived by averaging across cells that express gRNAs for the same target gene. **d)** CROP-seq data analysis identifies pathway signature genes and quantifies the effect of specific gRNAs on these signatures. **e)** CROP-seq uses a lentiviral construct in which the gRNA expression cassette is positioned within the 3’ long-terminal repeat (LTR), causing its duplication during lentiviral integration. It is expressed as part of two transcripts – a short RNA polymerase III transcript for genome editing and a long RNA polymerase II transcript that is poly-adenylated and detectable with Drop-seq. **f)** Lentiviral titers for LentiGuide-Puro (a standard CRISPR vector) and the CROPseq-Guide-Puro vector show that cloning into the 3’ LTR did not compromise the lentiviral function for gRNAs (in contrast, a 1,885 bp filler construct led to a strong reduction in viral titer). **g)** Genome editing efficiencies and indel signatures are similar between LentiGuide-Puro and CROPseq-Guide-Puro, based on next generation sequencing of PCR amplicons. **h)** CROP-seq can detect gRNAs from single-cell transcriptomes. **i)** Between 30% and 60% of single-cell transcriptomes are uniquely assigned to a gRNA, depending on the chosen threshold for the minimum number of genes detected in a cell. **j)** Performance statistics for single-cell RNA-seq across all CROP-seq experiments.

**Figure 2: CROP-seq analysis of T cell receptor signaling**

**a)** Experimental design of a single-cell CRISPR screen for the T cell receptor (TCR) pathway. Cas9-expressing Jurkat cells were infected with the CROPseq-Guide-Puro lentivirus containing a TCR-focused gRNA library at a low multiplicity of infection (MOI = 0.3). Lentivirus expressing cells were selected, and after 10 days the cell population was serum-starved, split, and either stimulated with anti-CD3/CD28 antibodies, or subjected to continued starvation. Cells were then subjected to single-cell RNA-seq. **b)** Fold change of gRNA abundance between the original plasmid pool and the CROP-seq experiments for naïve and TCR-stimulated cells. Values were normalized to the total of detected amplicons or assigned cells, respectively. **c)** Differential expression between TCR-stimulated cells and naïve Jurkat cells (histogram) and enriched pathways among the most differentially expressed genes (inlet). The x-axis shows the uncorrected posterior probability, and the chosen threshold for differentially expressed genes (red) corresponds to a false discovery rate of 5% using Holm-Bonferroni correction^25^. The five most enriched pathways based on the Enrichr tool^26^ tool are shown and sorted by their combined enrichment score (*p*-value * *z*-score). **d)** Heatmap of mean relative expression (column z-score) across the 569 pathway signature genes (columns), aggregated across cells that express gRNAs targeting the same gene (rows). **e)** Distribution of signature gene intensity across single cells (left) and number of cells (right) grouped by gRNA target gene. **(f)** Deviation of signature gene intensity for each gRNA target gene relative to control cells in TCR-stimulated Jurkat cells. **g)** CROP-seq results (signature gene intensity as shown in panel f) mapped to key components of the TCR pathway.

---

**Supplementary Figure 1.**
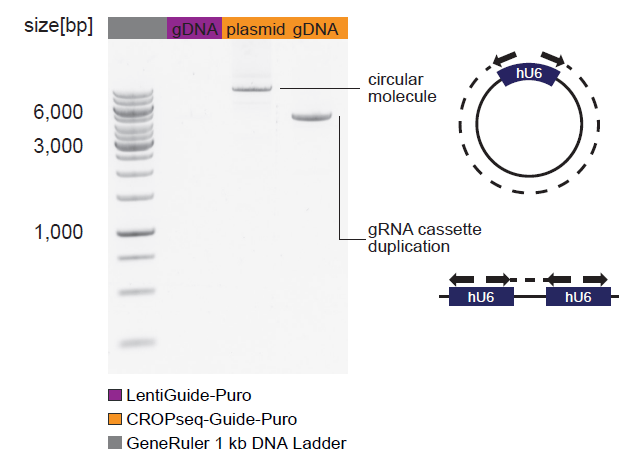
Validation of the hU6-gRNA cassette duplication in CROPseq-Guide-Puro infected cells. To validate the duplication of the hU6-gRNA cassette during lentiviral integration, we performed PCRs with primers binding to the hU6 promoter, but facing in opposite directions. Productive amplification can only occur when amplifying from a circular plasmid or after duplication of the cassette during lentiviral integration. As templates, we used gDNA from LentiGuide-Puro infected cells (lane 1, resulting in no amplification), a plasmid preparation of CROPseq-Guide-Puro (lane 2), or gDNA from CROPseq-Guide-Puro infected cells (lane 3).

**Supplementary Figure 2.**
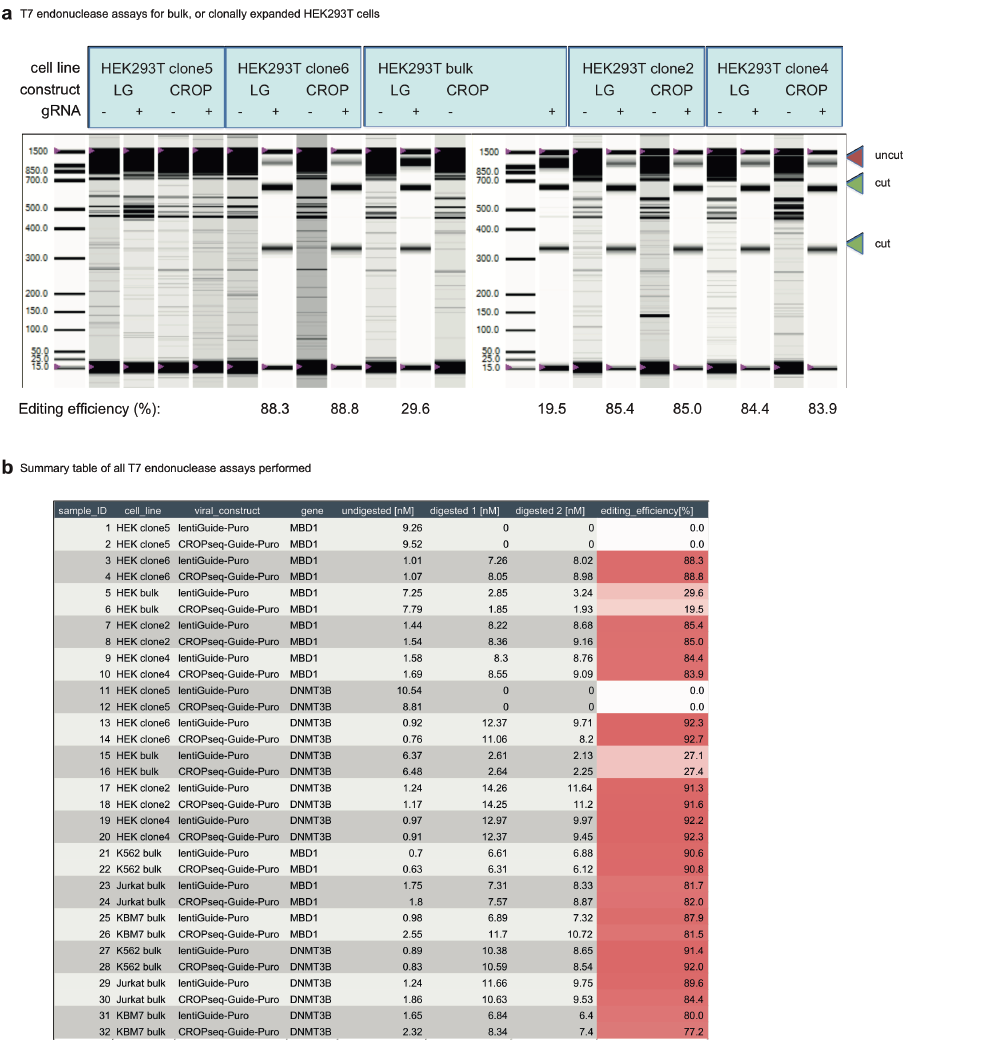
Similar genome editing efficiencies for LentiGuide-Puro and CROPseq-Guide-Puro based on the T7 endonuclease assay. **a)** Bulk and clonally expanded HEK293T cell lines were infected with LentiGuide-Puro (LG) or CROPseq-Guide-Puro (CROP) using vectors containing a gRNA targeting the MBD1 locus (+) or a different genomic locus (–). Genome editing efficiencies for MBD1 were estimated using the T7 endonuclease assay and highly similar between the two vectors. **b)** Table summarizing genome editing efficiencies for four cell lines (HEK293T, K562, Jurkat, KBM7) and two gRNAs (MBD1, DNMT3B).

**Supplementary Figure 3.**
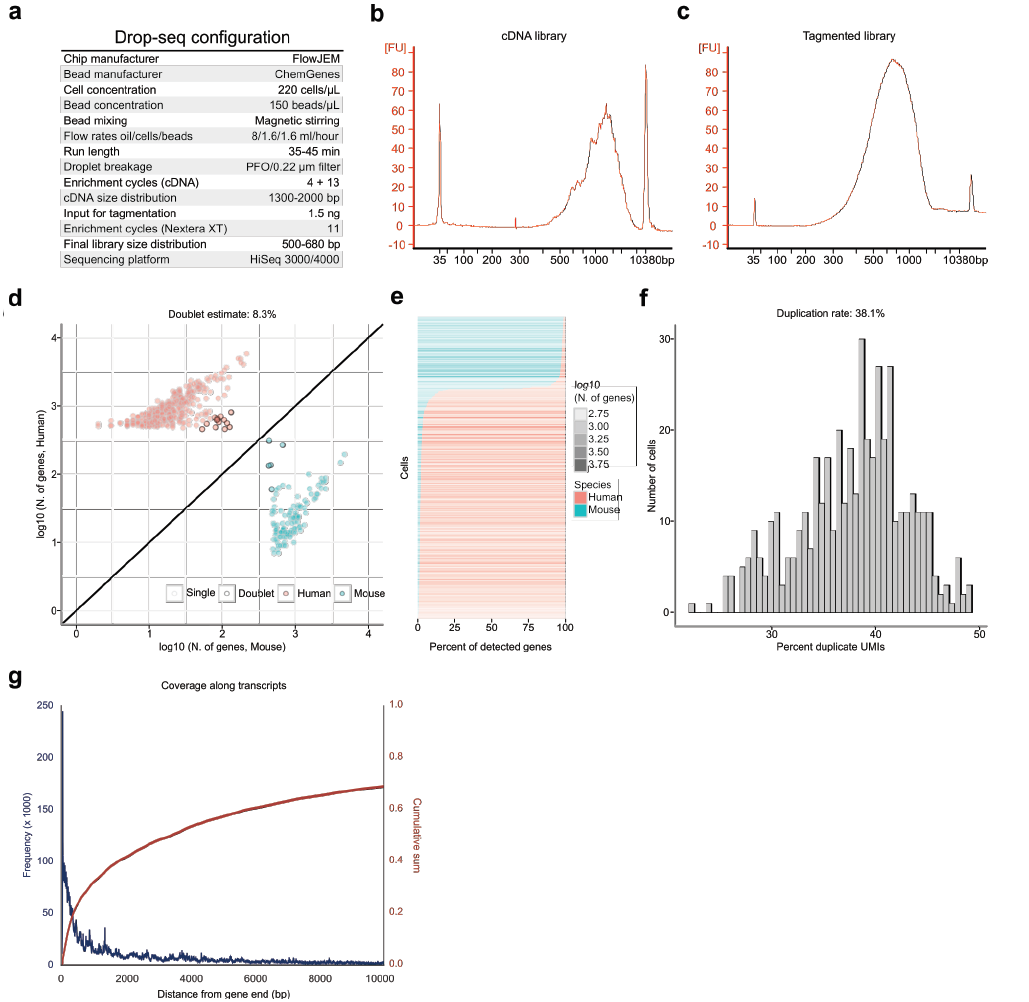
Configuration and validation of the Drop-seq method for single-cell transcriptome profiling. **a)** Setup of the Drop-seq assay used as a key component of CROP-seq. **b)** Bioanalyzer trace of a typical cDNA library from a Drop-seq run. **c)** Electropherogram of a tagmented library ready for sequencing. **d)** Doublet estimates based on a HEK293T (human) / 3T3 (mouse) mixing experiment. **e)** Percent of detected genes aligning to the human and mouse transcriptomes, shown for all cells with more than 500 genes. **f)** Duplicate rates estimated using unique molecular identifiers (UMIs). **g)** Distribution of the distance of mapping positions of transcriptome reads to the 3’end of gene models (blue line) and cumulative sum of the same (red line).

**Supplementary Figure 4.**
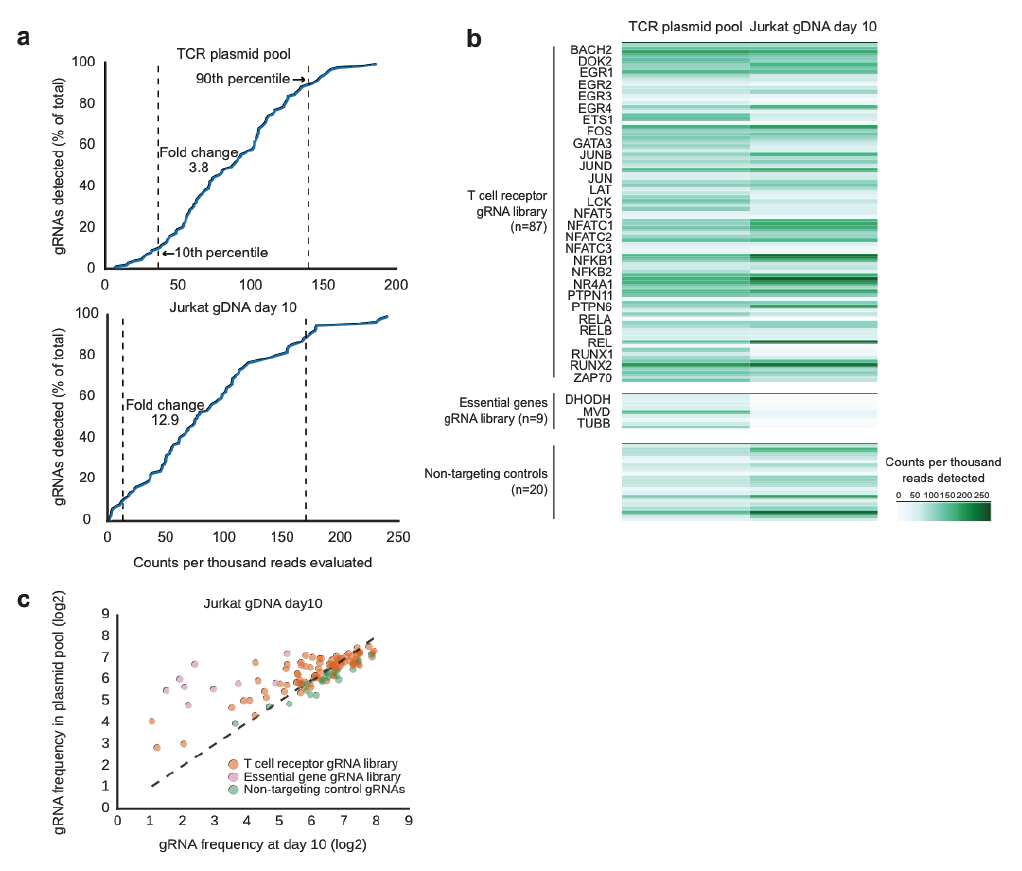
Validation of the TCR gRNA library complexity and gRNA dynamics in a pooled CRISPR screen. **a)** gRNA representation in the TCR gRNA library, assessed by amplicon sequencing of the plasmid pool (top) and the gDNA of Jurkat cells at day 10 post infection with CROPseq-Guide-Puro (bottom), both displayed as cumulative distribution plots. The fold change between the 10^th^ and 90^th^ percentile is highlighted as a measure of library imbalance, which expectedly increases upon infection. **b)** Abundance of each gRNA shown as a heatmap. **c)** Scatterplots of gRNA abundance from amplicon libraries at day 10 versus the original plasmid library. Frequencies of detected gRNAs have been normalized to the evaluated reads in each experiment.

**Supplementary Figure 5.**
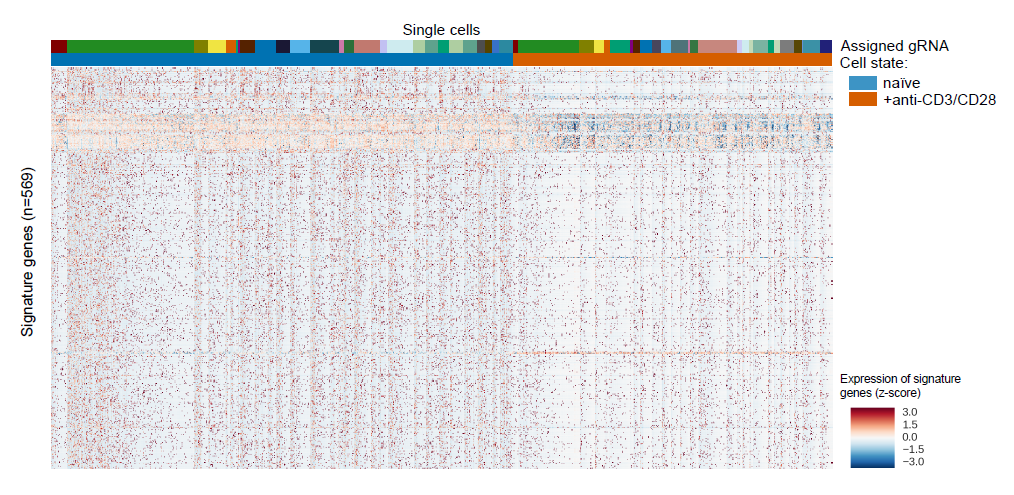
Single-cell transcriptome profiles for TCR stimulated Jurkat cells. Heatmap of single-cell expression data for TCR-specific signature genes (y-axis) shown for all cells with a uniquely assigned gRNA (x-axis). Genes were clustered hierarchically by the Euclidian distance, and cells were sorted by stimulation condition and gRNA assignment.

**Supplementary Figure 6.**
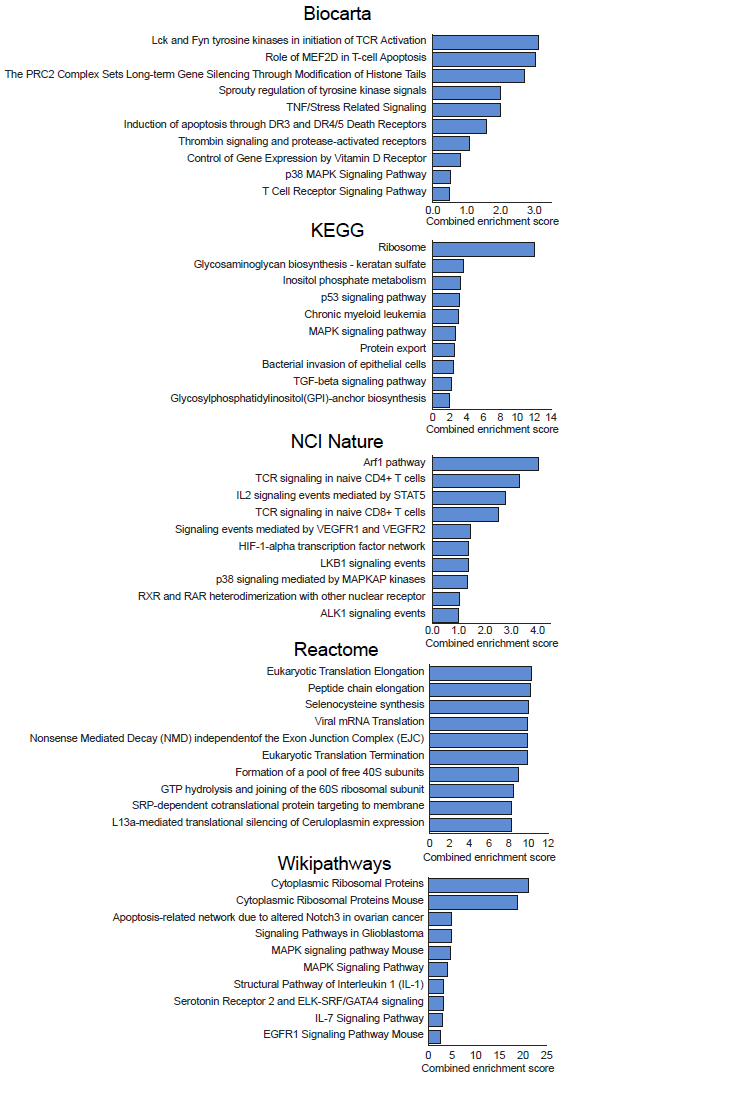
Pathway enrichment of TCR signature genes. Combined score [log(*p*-value) * *z*-score] of the enrichment of TCR signature genes across several gene set databases, as calculated by the Enrichr tool.

**Supplementary Figure 7.**
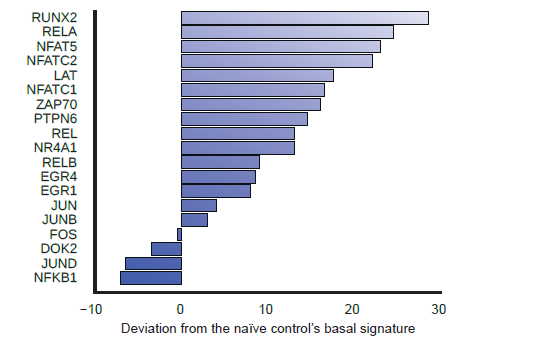
Target gene specific deviation from the basal Jurkat gene signature. Deviation of signature gene intensity for each gRNA target gene relative to control cells in unstimulated (naïve) Jurkat cells.

